# Plant responses to multiple antagonists are mediated by order of attack and phytohormone crosstalk

**DOI:** 10.1101/2021.02.17.431727

**Authors:** Saumik Basu, Robert E. Clark, Sayanta Bera, Clare L. Casteel, David W. Crowder

**Affiliations:** Department of Entomology, Washington State University, Pullman, WA, USA, 99164; School of Integrative Plant Science, Plant Pathology and Plant-Microbe Biology Section, Cornell University, Ithaca, NY, USA 14850

**Keywords:** disease ecology, plant-insect-pathogen interactions, plant defensive chemistry, plant nutrients, species interactions

## Abstract

Plants are often attacked by multiple antagonists, and traits of the attacking organisms, and their order of arrival onto hosts, may affect plant defenses. However, few studies have assessed how multiple antagonists, and varying attack order, affect plant defense or nutrition. To address this, we assessed defensive and nutritional responses of *Pisum sativum* plants after attack by a vector herbivore (*Acrythosiphon pisum*), a non-vector herbivore (*Sitona lineatus*), and a pathogen (*Pea enation mosaic virus*, PEMV). We show PEMV-infectious *A. pisum* induced several pathogen-specific plant defense signals, but these defenses were inhibited when *S. lineatus* was present in peas infected with PEMV. In contrast, feeding by *S. lineatus* induced anti-herbivore defense signals, but these defenses were enhanced by PEMV. *Sitona lineatus* also increased abundance of plant amino acids, but only when they attacked after PEMV-infectious *A. pisum*. Our results suggest that diverse communities of biotic antagonists alter defense and nutritional traits of plants through complex pathways that depend on the identity of attackers and their order of arrival onto hosts. Moreover, we show interactions among a group of biotic stressors can vary along a spectrum from antagonism to enhancement/synergism based on the identity and order of attackers, and these interactions are mediated by a multitude of phytohormone pathways.

## 1 INTRODUCTION

Plant hosts have defenses to counter attacks from antagonists such as herbivores and pathogens (Pandey, Ramegowda, & Senthil-Kumar, 2015; Miller, Costa Alves, & Van Sluys, 2017). For example, the jasmonic acid pathway often regulates plant defenses against herbivores, while the salicylic acid pathway often regulates defenses against pathogens (Koornneef & Pieterse, 2008; Thaler, Humphrey, & Whiteman, 2012). While many studies have assessed plant responses to particular antagonists, plants are often challenged by many stressors concurrently, and plant defenses can depend on the order in which antagonists arrive on plants (Thaler et al., 2012; Nejat & Mantri, 2017). For example, herbivores often limit plant defenses against pathogens when they arrive first on plants, but herbivores often have few impacts on plant defenses against pathogens when they arrive after pathogens on plants (Okada, Abe, & Arimura, 2015; Lin et al., 2019). In other contexts, certain organisms ‘prime’ pathways, promoting defense against subsequent organisms activating the same pathway, such that attack order may not matter (Mauch-Mani, Baccelli, Luna, & Flors, 2017; Ramírez-Carrasco, Martínez-Aguilar, & Alvarez-Venegas, 2017).

Biotic stressors may also alter the nutritional quality of plants by regulating amino acid metabolism (Casteel et al., 2014; Zhou, Lou, Tzin, & Jander, 2015), which can alter the feeding behavior and nutrient uptake by subsequent herbivores (Behmer, 2009; Zhu, Poelman, & Dicke, 2014). For example, the composition of free amino acids constitutively changes in leaves of soybean plants in response to soybean aphids (Chiozza, O’Neal, & MacIntosh, 2010). *Tomato yellow leaf curl virus* also alters the nutritional quality of tomato plants by affecting free amino acid levels in phloem, which alters the amino acid composition of whitefly (*Bemisia tabaci*) honeydew (Guo et al., 2019). However, few studies have explored how the diversity and identity of attacking organisms, and variation in the order of attack, affect nutritional traits of plants.

While there has been considerable research on the jasmonic and salicylic acid pathways, to understand complexities of plant defense it is necessary to assess how biotic antagonists mediate other signaling pathways (e.g., Lacerda, Vasconcelos, Pelegrini, & Grossi de Sa, 2014; Suzuki, 2016). Moreover, it is key to assess how changes in plant defense correlate with plant nutrients. For example, plants in low-nitrogen soil often adopt carbon-based defenses, while plants grown with fertilizer often accumulate more nitrogenous toxins (Cipollini, Walters, & Voelckel, 2017). Nitrogen in plants may also affect both pathogens and herbivores through synthesis of defensive metabolite, nitric oxide, and by nitrogen mobilization (War et al., 2012; Mur, Simpson, Kumari, Gupta, & Gupta, 2017). However, few studies have correlated effects of multiple biotic stressors on both plant chemical signaling and nutritional properties (Petek et al., 2014; Su et al., 2016).

We addressed these knowledge gaps by assessing the response of *Pisum sativum* plants to attack from a piercing-sucking vector herbivore, the pea aphid (*Acrythosiphon pisum*), a chewing non-vector herbivore, the pea leaf weevil (*Sitona lineatus*), and an aphid-borne pathogen, *Pea-enation mosaic virus* (PEMV). These organisms co-occur in ecosystems of eastern Washington and northern Idaho, USA, and interactions between them can affect plant traits and signaling pathways affecting insects and pathogens (Chisholm, Eigenbrode, Clark, Basu, & Crowder, 2019; Bera, Blundell, Liang, Crowder, & Casteel, 2020). However, the order in which herbivores and pathogens arrive on hosts, which varies across sites (Chisholm et al., 2019), may impact plant traits and defenses. To address this, we varied the diversity, identity, and order of attack among this community of biotic antagonists and assessed resulting changes in gene expression and phytohormones related to plant defense and nutrition. Our study revealed how plant responses to diverse stressors can mediate complex species interactions within a pathosystem.

## 2 MATERIALS AND METHODS

### 2.1 Study system

The Palouse region of eastern Washington and northern Idaho, USA, is home to many legumes including *P. sativum* (Black et al., 1998). In *P. sativum* fields, *S. lineatus*, a chewing herbivore, co-occurs with *A. pisum*, a phloem-feeding herbivore that can transmit pathogens such as PEMV (Chisholm et al., 2019). PEMV is one of several viruses that infects *P. sativum*, and this pathogen is obligately transmitted by aphids in a persistent manner (Chisholm et al., 2019).

*Sitona lineatus* adults overwinter outside of *P. sativum* fields and migrate into fields in late spring before *A. pisum* arrive (Cárcamo et al., 2018). After *S. lineatus* eggs hatch, larvae burrow into the soil to feed and pupate before emerging as adults in the summer (Cárcamo et al., 2018); these second-generation adults often occur on plants under attack from *A. pisum* and PEMV (Chisholm et al., 2019). Thus, *S. lineatus* attacks plants in the field both before and after *A. pisum* and PEMV. However, it is unknown if responses of *P. sativum* differ based on the number of stressors, and their order of attack. Moreover, molecular mechanisms that mediate interactions among these stressors are largely unknown (Chisholm et al., 2019; Bera et al., 2020).

To address these questions, we conducted greenhouse assays to assess interactions between *S. lineatus, A. pisum*, and PEMV on *P. sativum* plants, and molecular mechanisms affecting these interactions. First-generation adult *S. lineatus* for experiments were collected from commercial *P. sativum* fields, or wild patches of *Vicia villosa*, immediately prior to experiments. Colonies of infectious *A. pisum* with PEMV, and uninfectious *A. pisum*, were started from Palouse field-collected individuals (Chisholm et al., 2019) and were maintained on *P. sativum* plants in a greenhouse (21-24°C during day cycle, 16-18°C during dark cycle, 16:8 h light:dark).

### 2.2 Experimental design

We conducted a 3 × 2 greenhouse (21-24°C day cycle, 16-18°C dark cycle, 16:8 h light:dark) experiment that varied *S. lineatus, A. pisum*, and PEMV (Fig. 1). There were three *S. lineatus* treatments: (i) control: no adults prior to *A. pisum* treatments (none), (ii) two adults that fed for 48 h prior to *A. pisum* treatments (first), and (iii) two adults that fed for 48 h after *A. pisum* treatments (second). The two *A. pisum* treatments were: (i) sham: 10 5-d old uninfectious adults that fed for 48 h and (ii) PEMV: 10 5-d old PEMV-infectious adults that fed for 48 h. For treatments with *S. lineatus* first, they were removed by hand prior to *A. pisum* treatments; for treatments with *A. pisum* first, they were removed by aspirator prior to *S. lineatus* treatments. Treatments were conducted on individual *P. sativum* plants in mesh ‘bug dorms’ (0.6 × 0.6 × 0.6 m), with six replicates randomly assigned to each treatment in a factorial design (3 *S. lineatus* treatments × 2 *A. pisum* treatments). After insects were removed, plants were allowed to develop for 7 d before we harvested tissue to assess viral titer, gene expression, and nutrients. Tissue samples from the whole aboveground portion plants were collected and flash frozen in liquid nitrogen and stored in a -80°C freezer until processing. Viral titer samples confirmed that 100% of plants in the PEMV treatments became infected over the course of the experiment.

**Figure 1.**
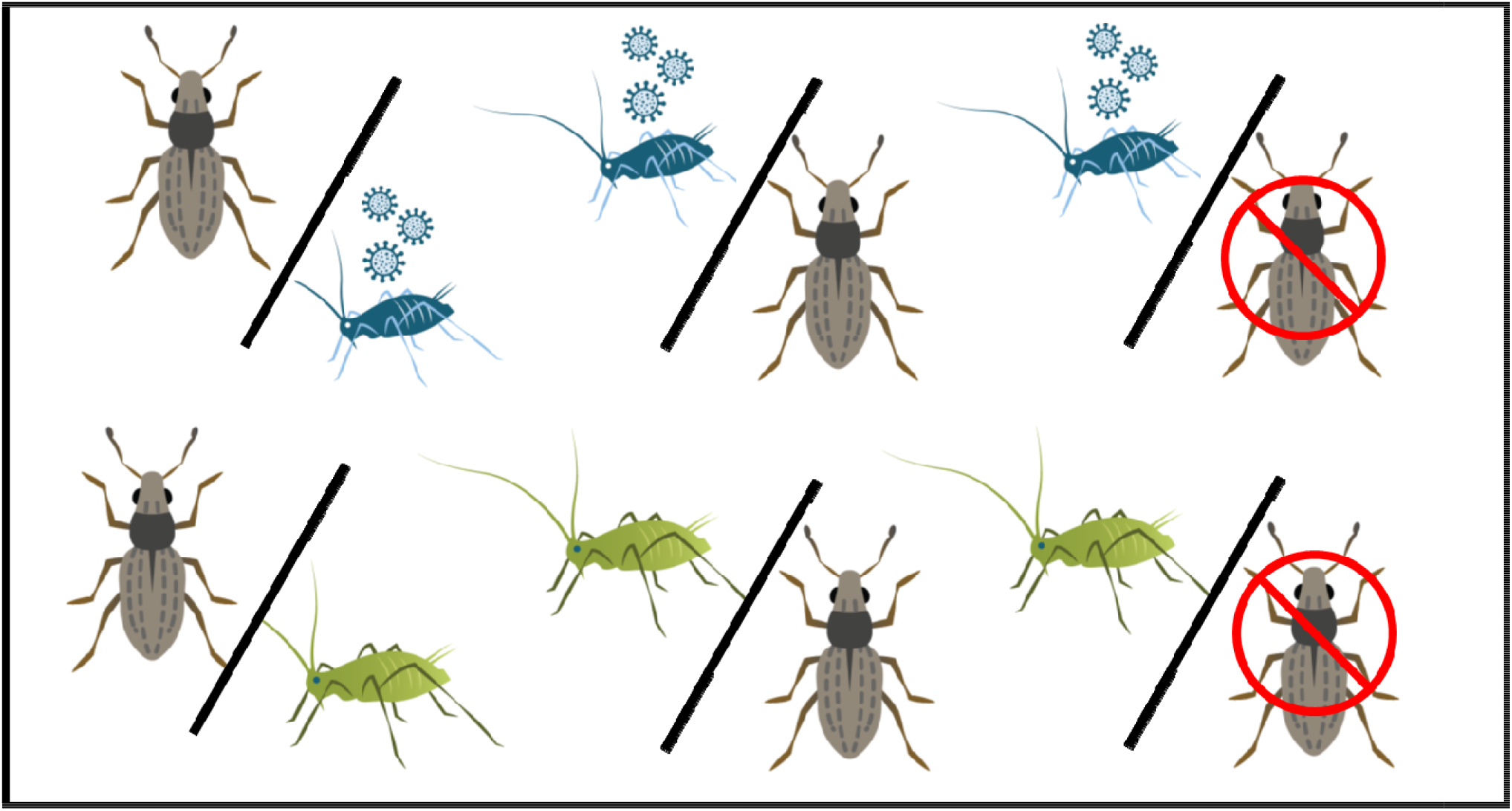
Schematic representation of 2 × 3 factorial design. Green-colored aphids indicate sham (non-infective) *A. pisum*, while blue-colored aphids indicate PEMV-infective *A. pisum*. Slashes indicate order of *S. lineatus* treatments (*S. lineatus* first, *A. pisum* first, or no *S. lineatus*).

### 2.3 Analysis of plant defense and biosynthetic genes

Plant tissue was processed using liquid nitrogen in sterilized mortars and pestles. Powdered tissue samples (50 to 100 mg) were used for RNA extraction with Promega SV total RNA isolation kits (Promega, Madison, WI). The quantity and quality of RNA was estimated on a NanoDrop1000 and agarose gel electrophoresis, respectively and 1 µg of total RNA from each sample was used for cDNA synthesis (Bio-Rad iScript cDNA Synthesis kits). Gene specific primers (Table S3) for qRT-PCR were designed using the IDT Primer Quest Tool. Each qRT-PCR reaction (10 µl) was set up containing 3 µl of ddH2O, 5 µl of iTaq Univer SYBR Green Supermix (Bio-Rad), 1 µl of specific primer mix (forward and reverse [concentration 10µM]), and 1 µl of diluted (1: 25) cDNA template. Reactions were set up in triplicates for each sample and ran on a CFX96 qRT-PCR machine (Biorad). The qRT-PCR program included an initial denaturation for 3 min at 95°C, followed by 40 denaturation cycles for 15 s at 95°C, annealing for 30 s at 60°C, and extension for 30 s at 72°C. For melting curve analysis, a dissociation step cycle (55°C for 10 s and then 0.5°C for 10 s until 95°C) was added. The comparative 2^−ΔΔCt^ method (Livak & Schmittgen, 2001; Kozera & Rapacz, 2013) was used to calculate the relative expression level of each gene, with β-tubulin as an endogenous control.

We assessed expression of seven genes associated with defense in peas. Gene sequences were obtained using accession numbers (available genes) or using Pea Marker Database (Kulaeva et al., 2017) and blast searching through the reference genome (Kreplak et al., 2019). Four genes were associated with plant hormone biosynthesis: (i) *Isochorismate synthase1* (*ICS1*) (salicylic acid), (ii) *Lipoxygenase 2* (*LOX2*) (jasmonic acid), (iii) *Aldehyde oxidase 3* (*AO3*) (abscisic acid), and (iv) *Gibberellin 2-oxidase (GA2ox*) (gibberellic acid). *ICS1* converts chorismate to isochorismate, a precursor of salicylic acid biosynthesis (Seguel et al., 2018), while *LOX2* is a precursor to jasmonic acid biosynthesis (Wasternack & Hause, 2013). *AO3* catalyzes abscisic acid biosynthesis by oxidizing abscisic aldehyde, and *GA2OX* catalyzes bioactive giberrelic acids or their immediate precursors to inactive forms (Zdunek-Zastocka & Sobczak, 2013; Serova, Tsyganova, Tikhonovich, & Tsyganov, 2019; He et al., 2019). All of these gene transcripts can affect plant defense and plant-microbe interactions (Lee et al., 2012; Yergaliyev et al., 2016).

The three additional genes examined were associated with defense response transcripts that occur downstream from hormone induction. One of these genes, *Pathogenesis-related protein 1* (*PR1*) affects systemic acquired resistance-mediated defense signaling and occurs downstream in the salicylic acid pathway (Fondevilla, Küster, Krajinski, Cubero, & Rubiales, 2011; Miranda et al., 2017). The second defense response transcript was an antimicrobial defensin peptide called *Disease resistance response* gene *(DRR230)*, which has been reported to provide resistance in peas against various pathogens (Lacerda et al., 2014; Selim, Sanssené, Rossard, & Courtois, 2017). The third defense response transcript assessed was *Lectin (PsLectin)*. Plant lectins are a group of carbohydrate binding proteins, and *Lectin* genes can be induced by salicylic acid, jasmonic acid, and herbivores to stimulate phytoalexin and pistatin production in peas (Fondevilla et al., 2011; Armijo et al., 2013; Macedo, Oliveira, & Oliveira, 2015).

### 2.4 Measurement of plant phytohormones

Plant tissue samples were assessed for three phytohormones: jasmonic acid, salicylic acid, and abscisic acid following procedures of Patton, Bak, Sayre, Heck, & Casteel (2019). Briefly, tissue samples were first flash frozen in liquid nitrogen before being lyophilized and weighed. Hormones were extracted in iso-propanol:H_2_O:HCL_1MOL_ (2:1:0.005) with 100 μl of internal standard solution (1000 pg of each). Samples were evaporated to dryness, resuspended in 100 μl of MeOH, filtered, and 10 µl of each sample was injected into an Agilent Technologies 6420 triple quad liquid chromatography-tandem mass spectrometry instrument (Agilent, Santa Clara, CA). A Zorbax Extend-C18 column 3.0 × 150mm (Agilent, Santa Clara, CA) was used with 0.1% formic acid in water (A) and 0.1% (v/v) formic acid in acetonitrile (B) at a flow rate of 600 mL min^−1^. The gradient used was 0–1 min, 20% B; 1–10 min, linear gradient to 100% B; 10-13 min, 100% A. Retention times were: jasmonic acid (D5) standard (5.740 min), jasmonic acid (5.744 min), salicylic acid D4 standard (4.677 min), salicylic acid (4.720 min).

### 2.5 Analysis of plant nutritional components

For amino acid analysis, leaf tissue was lyophilized, weighed, and extracted with 20mM of HCL (Patton et al., 2019). Derivation was done using AccQTag reagents following the manufacturer’s instructions (Waters, Shinagawa-ku, Tokyo), and derivatised samples (10 μl) were then injected. Ground tissue was extracted with 100 μl of 20 mM HCl, centrifuged, and the supernatant was saved. Amino acids were derivatized using AccQ-Fluor reagent kits (Waters, Milford, MA), with L-Norleucine as an internal standard. 10 μl from each sample were injected with an Agilent 1260 Infinity pump with a Nova-Pak C18 column and fluorescence detector, and Agilent Chemstation software for data recording. Amino acid derivatives were detected with an excitation wavelength of 250 nm and an emission wavelength of 395 nm. Peak areas were compared to a standard curve made from a serial dilution of amino acid standards (Sigma-Aldrich, St. Louis, MO). injected into a Agilent 1260 Infinity HPLC (Agilent, Santa Clara, CA) with a Nova-Pak C18 column (Casteel et al., 2014). Solvent A, AccQ-Tag Eluent A, was premixed from Waters; Solvent B was acetonitrile:water (60:40). The gradient used was 0–0.01 min, 100% A; 0.01–0.5 min, linear gradient to 3% B; 0.5–12 min, linear gradient to 5% B; 12–15 min, linear gradient to 8% B; 15–45 min, 35% B; 45–49 min, linear gradient to 35% B; 50–60 min, 100% B. The flow rate was 1.0 ml min^−1^. Amino acid derivatives were measured with an Agilent fluorescence detector with an excitation wavelength of 250 nm and an emission wavelength of 395 nm. For concentration calculations, standard curves were generated for each amino acid using dilutions of the standard.

### 2.6 Data analysis

To evaluate effects of our treatments on host-plant defenses and host-plant quality, we ran a series of multivariate models using R ver. 3.5.2 (R Working Group, 2018). First, gene expression was evaluated with *ICS1, LOX2, GA2ox, AO3, PR1, DRR230*, and *PsLectin* as the responses, with MANOVA to assess treatment effects on relative gene expression (2^−ΔΔCt^) based on cycle threshold values for each observed gene transcript. Estimated marginal mean of Ct values, and standard error of the mean, were generated using the emmeans package in R (Lenth, 2016). The methodology for 2^− ΔΔCt^ followed modified recommendations from Rao, Huang, Zhou, & Lin (2013) and Kozera & Rapacz (2013), using housekeeping gene *β-tubulin* to normalize expression and a sham aphid (non-infective Pea aphid and no weevil addition) treatment as a control.

Hormone levels in plants were evaluated using MANOVA, with salicylic acid, jasmonic acid, and abscisic acid as responses (3 variables). Total amino acid content was evaluated using a generalized linear model (GLM) with total concentration among all amino acids as the response. All models assessed treatment effects, using *S. lineatus* addition, *A. pisum* infection status, and their interaction as predictors. Finally, changes in the amino acid profile was evaluated using non-metric multidimensional scaling (NMDS) with the vegan package (Oksanen et al., 2019) following Ceulemans, Hulsmans, Ende, & Honnay (2017).

## 3 RESULTS

### 3.1 Effects of multiple antagonists and attack order on plant gene transcripts

Transcription of plant genes associated with hormone biosynthesis across four pathways:

(i) salicylic acid (*ICS1*), (ii) jasmonic acid (*LOX2*), (iii) abscisic acid (*AO3)*, and (iv) gibberellic acid (*GA2ox*), were induced by PEMV when *S. lineatus* was not present (Fig. 2A-D, Table S1, *Pillai =* 0.942, *P =* 0.002). However, there was no induction of any of these biosynthesis genes in response to PEMV when *S. lineatus* was present, indicating that *S. lineatus* inhibited plant defense against PEMV (Table S1, A xW interaction, *Pillai* = 1.509, *P* = 0.021, Fig. 2). In both the presence and absence of PEMV, *S. lineatus* induced transcription of *LOX2*, but *S. lineatus* did not directly modify the expression of ICS1, AO3 or GA2ox (Fig. 2B-D, Table S1). When PEMV-infectious *A. pisum* attacked following *S. lineatus*, there was greater induction of LOX2 compared to when *S. lineatus* attacked alone (Table S1, A x W interaction, *Pillai* = 1.509, *P* = 0.021, Fig. 2). In contrast to the antagonism exerted by *S. lineatus* on the response to PEMV, this represents enhancement of plant defense when PEMV infection followed attack by *S. lineatus*.

**Figure 2.**
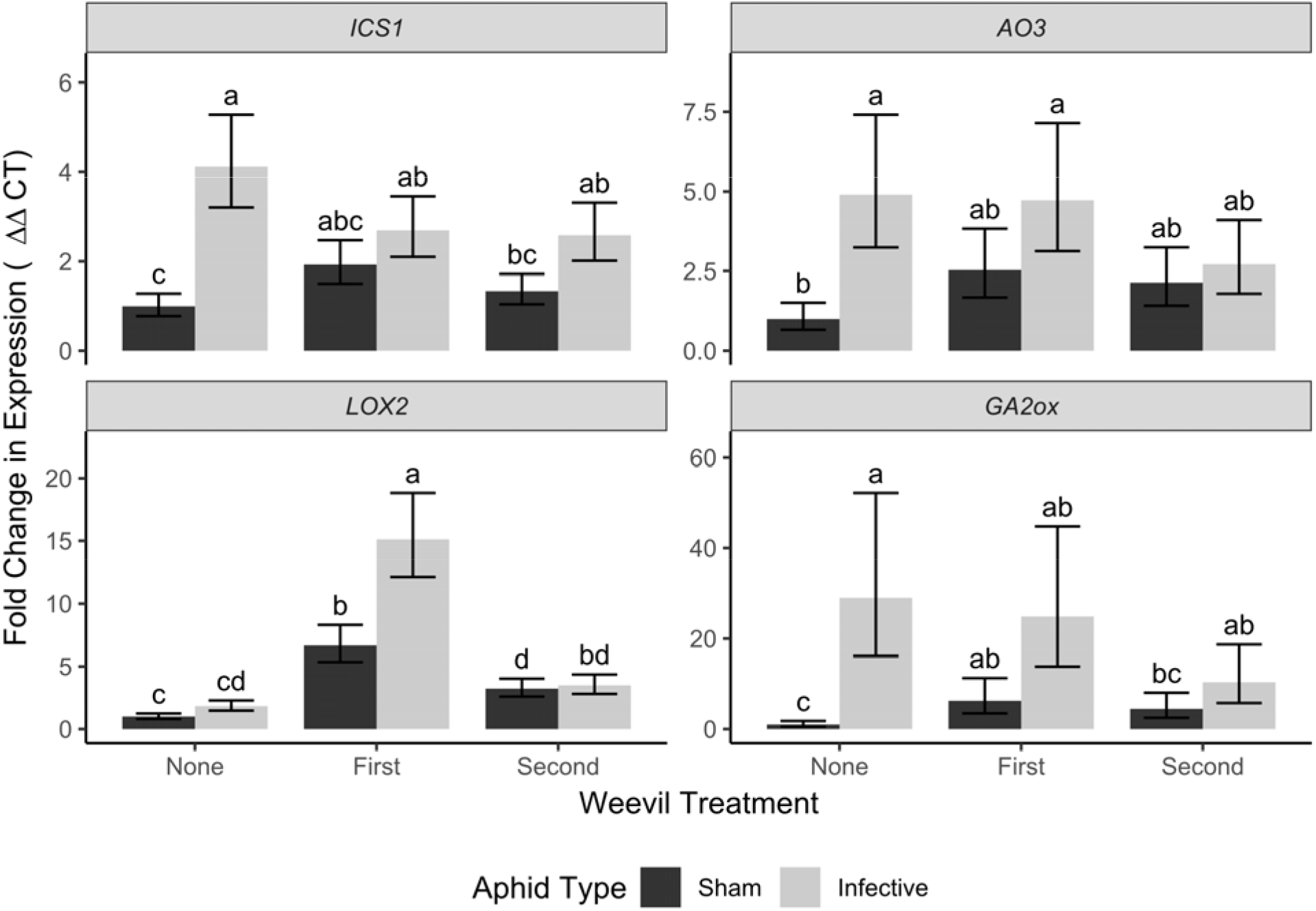
Relative transcript accumulation of plant hormone biosynthesis genes associated with four hormonal signaling pathways: (A) *ICS1* (salicylic acid), (B) *LOX2* (jasmonic acid), (C) *AOX3* (abscisic acid), and (D) *GA2ox* (gibberellic acid) following attack with various combinations of *S. lineatus, A. pisum*, and PEMV. Within each panel, bars separated by a different letter were significant different based on MANOVA (Tukey HSD, α = 0.05). Bar height and error bars indicate marginal mean and standard error of the regression coefficient for each respective treatment.

All three defense response transcripts (*PR1, DDR230, PsLectin*) were induced by PEMV when *S. lineatus* was not present (Fig. 3); similarly, each transcript was induced by *S. lineatus* when PEMV was not present (Fig. 3, Table S1; A ⍰ W interaction, *F =* 2.64, *P =* 0.111). When *S. lineatus* attacked second, the expression level of *PR1 and Lectin* did not change compared to when weevils were absent. The effects of PEMV on the transcripts was modified by the presence of *S. lineatus* and attack order. While *DDR230* was induced by PEMV (Table S1, *F* = 47.181, *P* < 0.001), this effect diminished when *S. lineatus* was present after PEMV (Fig. 3B). Similarly, the effects of PEMV on *PR1* were inhibited when *S. lineatus* attacked second (Fig. 3), whereas that the induction of lectin by PEMV was not altered by *S. lineatus* in either order (Fig. 3).2370

**Figure 3.**
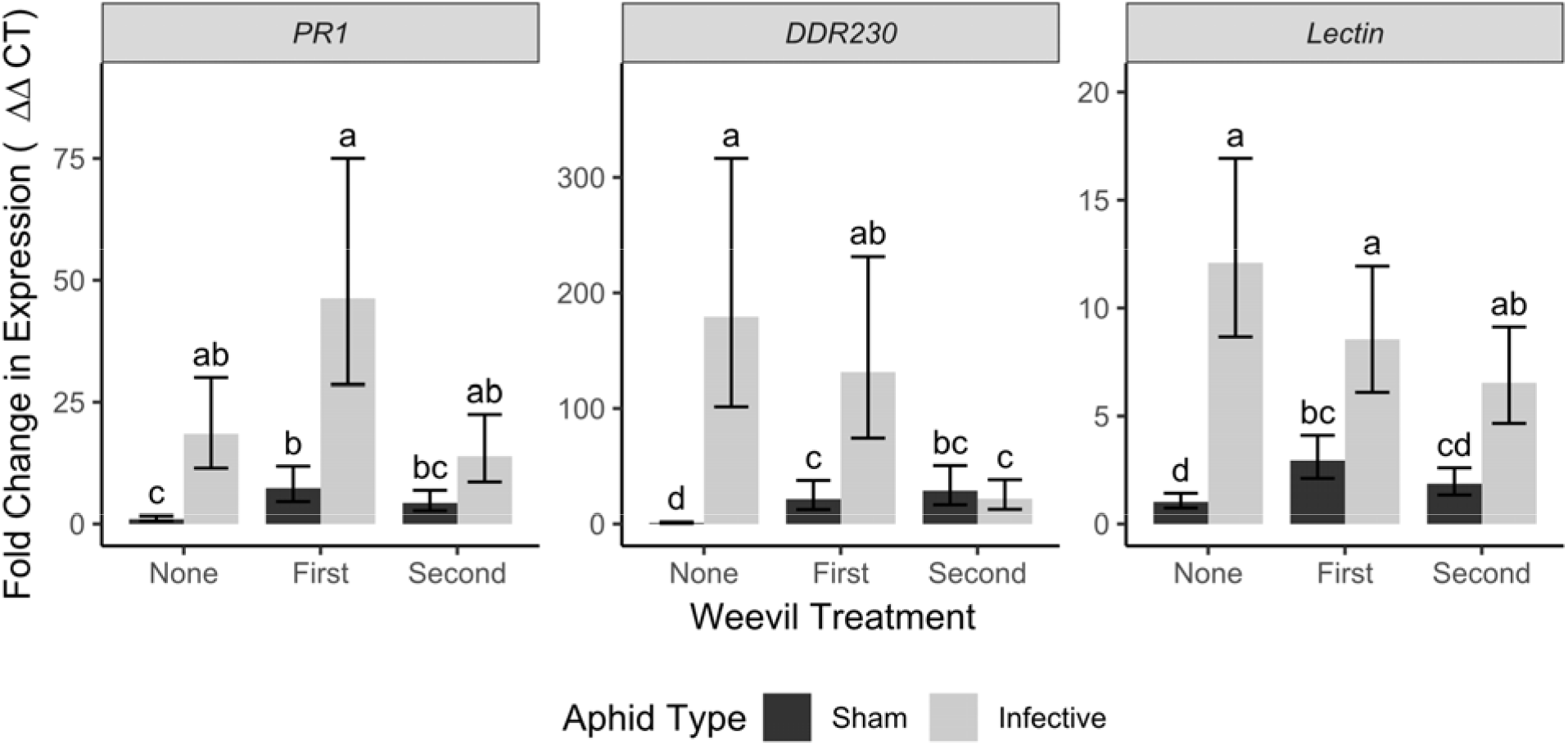
Relative transcript accumulation of plant defense response transcripts: (A) *PR1*, (B) *DDR230*, and (C) *PsLectin* following attack with various combinations of *S. lineatus, A. pisum*, and PEMV. Within each panel, bars separated by a different letter were significant different based on MANOVA (Tukey HSD, α = 0.05). Bar height and error bars indicate marginal mean and standard error of the regression coefficient for each respective treatment.

### 3.2 Effects of multiple antagonists and attack order on plant phytohormones

We observed variation in phytohormones in response to *A. pisum* (Table S2, *Pillai* = 0.95, *P* < 0.001) and *S. lineatus* (Table S2, *Pillai* = 1.195, P < 0.001). PEMV-infectious *A. pisum* strongly induced salicylic acid (Table S2, *F =* 254.2, *P* < 0.001), but this was inhibited when *S. lineatus* attacked after PEMV (Fig. 4A, Tukey HSD). PEMV did not affect jasmonic acid (Table S2, *F* = 0.97, P = 0.34), but the order of *S. lineatus* did (Table S2, *F* = 5.30, *P* = 0.018). Both *S. lineatus* (Table S2, *F* = 4.10, P = 0.037) and infectious *A. pisum* induced abscisic acid (Table S2, *F* = 9.96, P = 0.006) and this effect was contingent on the attack order (Table S2, A ⍰ W, *F =* 4.32, *P =* 0.032, Fig. 4, Tukey HSD). Jasmonic acid levels were suppressed by *S. lineatus* when attacking prior to non-infectious sham *A. pisum*, but not on plants already attacked by PEMV (Fig. 4B, Tukey HSD).

**Figure 4.**
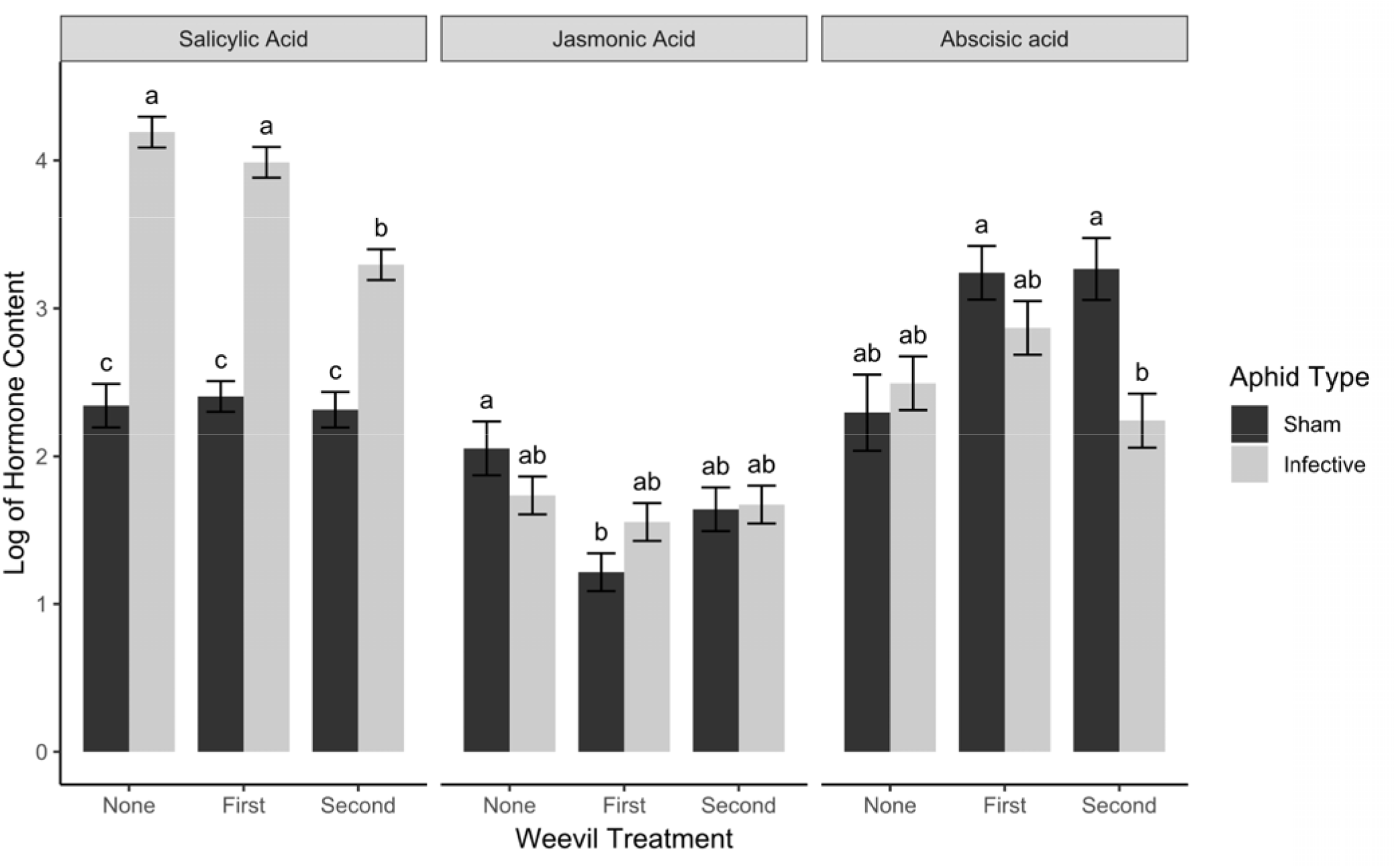
(A) Salicylic acid, (B) jasmonic acid, and (C) abscisic acid phytohormone levels in *P. sativum* plants following attack with various combinations of *S. lineatus, A. pisum*, and PEMV. Within each panel, bars not connected by the same letter are significantly different (Tukey HSD, α = 0.05). Bar height and error bars indicate marginal mean and standard error of the regression coefficient for each respective treatment.

### 3.3 Effects of multiple antagonists and attack order on plant nutrients

Feeding by *S. lineatus* increased the total amino acid levels (GLM, χ2 = 9.19, *P =* 0.01, Fig 5), but PEMV-infectious *A. pisum* did not (GLM, χ^2^ = 0.044, *P =* 0.83), and this effect was not modified depending on attack order (GLM, A ⍰ W interaction, χ^2^ = 0.24, *P =* 0.63). Non-metric multidimensional scaling (NMDS) analysis of amino acid composition also showed that changes to amino acid availability was most different among treatments for alanine, arginine, lysine, and glycine (Ordination plot, Fig S1).

**Figure 5.**
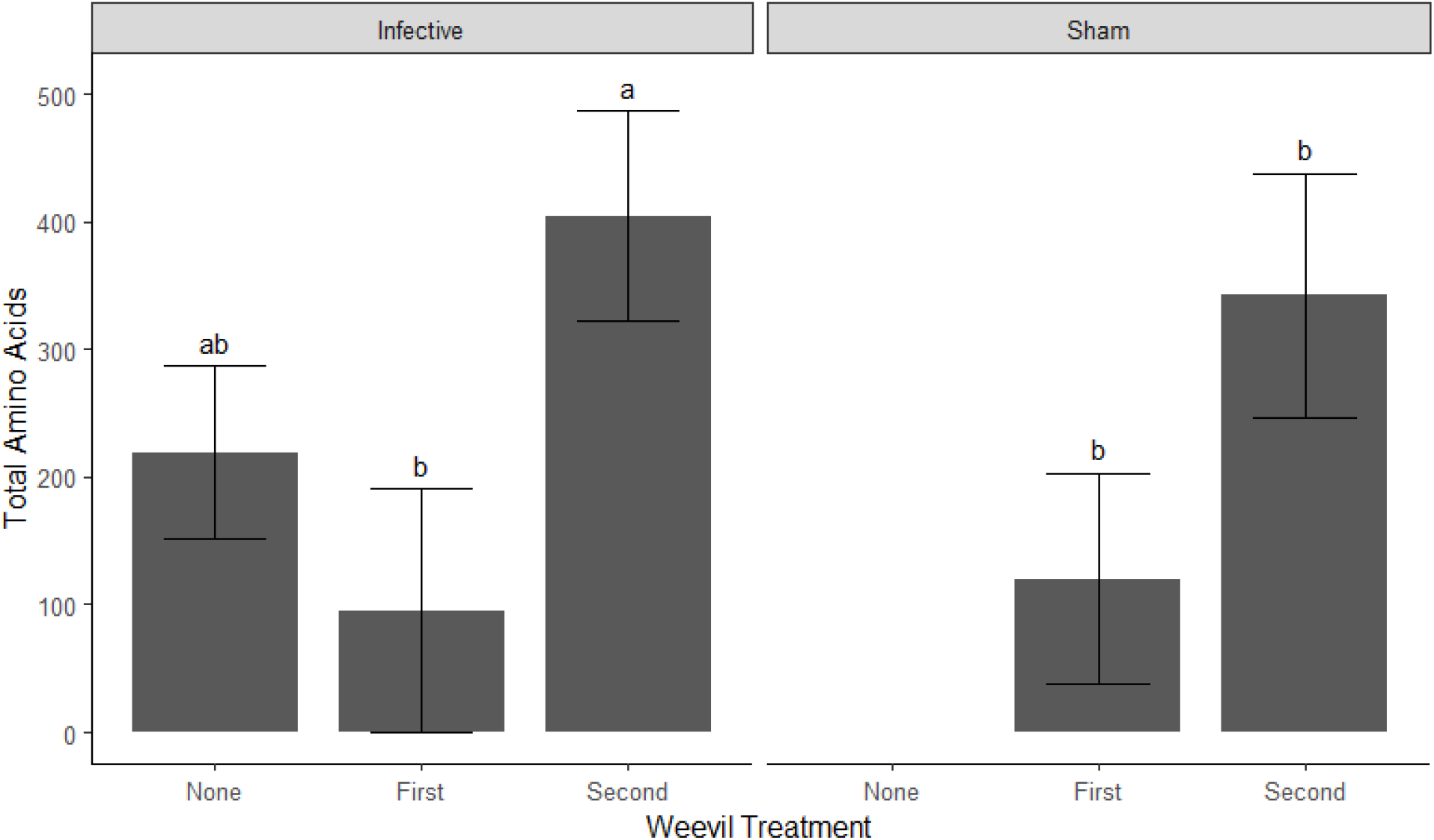
Nutritional analysis (total amino acid) in *P. sativum* following attack with various combinations of *S. lineatus, A. pisum*, and PEMV. *S. lineatus* increase total amino acid concentration in plants (GLM, χ2 = 9.194, *P =* 0.01). There was no “sham-none” treatment combination so that could not be estimated. Within each panel, bars not connected by the same letter are significantly different (Tukey HSD, α = 0.05). Bar height and error bars indicate marginal mean and standard error of the regression coefficient for each respective treatment.

## 5 DISCUSSION

Assessing interactions between biotic stressors and plants in food webs is critical to understand the dynamics of these interactions. Plant responses to stressors are often specific to the attacker, and both phytochemical responses and plant nutritional status can affect susceptibility to specific stressors (van Geem, Gols, Raaijmakers, & Harvey, 2016; Shikano, 2017). We show plants responded in complex ways to unique biotic stressors, including a piercing-sucking herbivore (*A. pisum*), a chewing herbivore (*S. lineatus*), and a virus (PEMV). Our study is among the first to assess how the order of attack, and diversity of stressors, mediate the defensive responses of plants and plant nutritional status (see also Vos, Moritz, Pieterse, & Van Wees, 2015). Our results show that plants traits varied in response to the type of attacker, the number of stressors, and their order of arrival. Moreover, we show that assessment of multiple gene transcripts, phytohormones, and plant nutrients provides a more comprehensive perspective on mechanisms driving plant-insect-pathogen interactions than any isolated response.

We found PEMV caused broad defensive responses in *P. sativum* by inducing specific gene transcripts and phytohormones (Figs. 2, 3 & 4). Biotropic pathogens such as PEMV are known to activate salicylic acid signaling (Singh, Swain, Singh, & Nandi, 2018; Chisholm, Sertsuvalkul, Casteel, & Crowder, 2018). This was reflected by increased expression of the *ICS1* biosynthesis gene, increased salicylic acid hormone levels, and increased expression of the downstream defensive transcript *PR1* when PEMV was present. However, effects of PEMV were not limited to salicylic acid, as PEMV induced gene transcripts often associated with biosynthesis of abscisic acid *(AO3*), and giberrellic acid (*GA2ox*), while also affecting defense response genes that occur downstream from induction of these hormones (*PR1, DRR230, PsLectin*). However, PEMV alone can not induce gene transcript associated with biosynthesis of JA (*LOX2*), but PEMV infection after weevil herbivory induced *LOX2* accumulation. Similar results in *P. sativum* have been observed in response to fungal infection by *Mycosphaerella pinodes* and *Phoma koolunga*, where infections induce defense related genes across multiple signaling pathways (Fondevilla et al., 2011; Tran, You, & Barbetti, 2018). However, increased expression of *LOX2* and *AO3* gene transcripts (Fig. 2) were not reflected by increased levels of jasmonic acid or abscisic acid (Fig. 4). This suggests that measuring phytohormones, or gene transcripts, in isolation may fail to reveal more complex pathways by which plants respond to stress (Kazan & Lyons, 2014).

While PEMV had broad effects on plants, *S. lineatus* attenuated these responses. When *S. lineatus* was present, before or after PEMV, the expression of three out of four biosynthesis gene transcripts (with the exception of *LOX2*) in response to infectious aphids were comparatively weaker (Fig. 2). Increased expression of *LOX2* may be due to *S. lineatus* inducing expression of defensive transcripts associated with JA-mediated chewing herbivore attack and the effect was further enhanced by PEMV infection after *S. lineatus* feeding (Fig. 2). Effects of PEMV on plant defense genes (*PR1, DDR230, PsLectin*) were also affected by *S. lineatus*, but varied with attack order. Overall, in this study the order of attack seemed to have stronger effects on downstream plant defense gene transcripts than on hormone biosynthesis gene transcripts.

While *S. lineatus* increased expression of two (*LOX2* and *GA2ox*) of the four biosynthesis gene transcripts studied when PEMV was not present, expression of *LOX2* was enhanced when PEMV was also present (Fig. 2). In contrast, PEMV caused decreased expression of two genes (*PR1, DRR230*) that were induced by *S. lineatus* was present alone (Fig. 2). For plants attacked first by either PEMV or *S. lineatus*, we observed the strongest evidence for mutual antagonism at the gene transcript level rather than for phytohormones (Figs. 2-4). Our study shows interactions among a set of stressors can vary based on attack order. Here, we found that *S. lineatus* feeding following PEMV infection inhibited plant defense (mutual antagonism), while *LOX2* expression was enhanced when PEMV infection followed *S. lineatus* feeding (synergistic effect). While the first effect is in line with studies showing “mutual antagonism” between chewing herbivores and biotropic pathogens (Thaler, Agrawal, & Halitschke, 2010; Vos et al., 2015), our study suggests the order of attack can lead to variation along a spectrum from antagonism to enhancement.

Our results provide evidence that the order of arrival of biotic stressors on plants can play a crucial role in determining plants’ response to these attackers. While mutual antagonism between *S. lineatus* and PEMV was common, for some genes these effects only occurred when *S. lineatus* attacked first, and for others when *S. lineatus* attacked second (Figs. 2-4). Mutual antagonism has most often been studied as effects of a prior attacker affecting a subsequent attacker, such as when a herbivore alters gene activation or phytohormones in ways that attenuate performance of subsequent attackers (Kessler & Halitschke, 2007; Erb, Robert, Hibbard, & Turlings, 2011; Stam, Mantelin, McLellan, & Thilliez, 2014; Huang et al., 2017). However, our results suggest that a second attacker may also mitigate defensive responses against the first attacker in ways that might affect plant defense and propagation of pathogens. For example, we show that plants infected by PEMV had decreased defenses when subsequently attacked by *S. lineatus* (Fig. 3), which should promote PEMV replication. Moreover, our results suggest that, PEMV infection induces pathogen defense and *S. lineatus* inhibits that if they appear on plants after the infection has been established. This may be more strongly expressed as variation in gene transcripts rather than hormone levels, a result that has similarly been seen in *Arabidopsis* in response to pathogen infection (Anderson et al., 2004).

Mutual antagonism in plant signaling pathways has most commonly been examined in regard to tradeoffs between jasmonic acid and salicylic acid. Our results show these tradeoffs extend to other signaling pathways. For example, jasmonic acid exhibits antagonism with abscisic acid in *Arabidopsis* following attack from *Fusarium oxysporum* (Anderson et al., 2004). Mutual antagonism between jasmonic acid and gibberellic acid, and jasmonic acid and abscisic acid, have also been reported (Yang, Yang, & He, 2013; Okada et al., 2015; Liu & Hou, 2018). For example, jasmonic acid facilitates defense over growth by repressing degradation of DELLA protein in rice and *Arabidopsis*, but elevated DELLA proteins interfere with the gibberellic acid pathway by binding to growth promoting transcription factors associated with gibberellic acid signaling (Yang et al., 2012, 2013; Okada et al., 2015). Antagonistic relationships between giberellic acid and abscisic acid have also been reported in both mono and dicot plants and regulated by various transcription factor regulators in response to diverse environmental cues (Liu & Hou, 2018). However, antagonisms between salicylic acid and abscisic acid may actually lead to synergism between jasmonic acid and abscisic acid, where elevated abscisic acid levels following infection with *Pseudomonas syringae* induce jasmonic acid in *Arabidopsis*, which in turn limits the levels of salicylic acid (Fan, Hill, Crooks, Doerner, & Lamb, 2009). Overall, these results suggest that a broad examination of genes and hormones are needed to elucidate pathways underlying plant-insect-pathogen interactions in *P. sativum* and other plants.

Our results suggest mutual antagonism may also occur among defense gene transcripts that are associated with a single signaling pathway. For example, the induction of *PR1*, a salicylic acid-responsive gene, was mitigated by *S. lineatus* attack after PEMV infection, as may be expected with antagonism between jasmonic acid and salicylic acid. However, the expression of *ICS1*, another gene associated with the biosynthesis of salicylic acid, was not responsive to *S. lineatus*. This has been seen in other studies where *ICS1* was not induced by caterpillar feeding although other genes associated with salicylic acid were (Onkokesung, Reichelt, van Doorn, Schuurink, & Dicke 2016). These results suggest that a plant’s response to multiple stressors is unlikely to result from simple crosstalk but rather from interactions among multiple signaling pathways that may exhibit complex responses.

In addition to affecting plant gene expression and phytohormones, plant pathogens such as viruses can also alter nutritional quality of their host plants in ways that affect vectors (Mauck, BosquelJPérez, Eigenbrode, Moraes, & Mescher, 2012; Wang, Senthil-Kumar, Ryu, Kang, & Mysore, 2012; Patton et al., 2019). Similarly, non-vector herbivores may strongly affect the quantity and quality of plant nutrients (Ángeles-López, Rivera-Bustamante, & Heil, 2016). For example, *pepper golden mosaic virus* (PGMV) infection in *Capsicum annuum* increased levels of the amino acids proline, tyrosine, valine but decreased levels of histidine and alanine. In the same system, the greenhouse whitefly, *Trialeurodes vaporarioum*, reversed the levels of these amino acids (Ángeles-López et al., 2016). Arrival of *S. lineatus* before PEMV infection suppressed the amount of total amino acids in peas, while enhanced amino acid level was detected if *S. lineatus* damaged peas after PEMV infected was established. This suggests the intriguing possibility that antagonism between a pathogen and non-vector herbivore can occur at the level of amino acid production in plants.

Overall, our study provides example of complex interactions between a vector-borne plant pathogen and a non-vector herbivore that varies from antagonism to enhancement and manifest as changes in plant gene transcripts, phytohormones levels, and plant nutrients. However, we show that assessing the order of attack is necessary to best understand the complexity and mechanisms of plant-insect-pathogen interactions. Moreover, our study suggests complete pathways must be characterized as differences are evident even when a few transcripts and metabolites are analyzed., often measured with associated gene transcripts (Bedini, Mercy, Schneider, Franken, & Lucic-Mercy, 2018; Ángeles-López el al., 2016; Shi et al., 2019), may fail to capture mechanisms by which plants interact with multiple stressors. Our results demonstrate both the order of arrival, and the diversity of interactions, determine plant responses to stress through the combined action of defense gene activation, phytohormone accumulation, and modification of plant nutrients. Characterizing the pathways by which plants respond to single and multiple stressors, with varying attack order, can shed light on the mechanisms that shape food web interactions among plants, herbivores, and pathogens.

## ACKNOWLEDGEMENTS

We thank the many undergraduates who helped in various experiments, data collection, and O. Hill for assistance with data analysis. This research was supported by USDA□NIFA Grants 2016□67011□24693, 2017-67013-26537, and Hatch project 1014754.

## DATA ACCESSIBILITY

Data are publicly available from Figshare: https://doi.org/10.6084/m9.figshare.14227097.v1

## AUTHOR CONTRIBUTIONS

S.B. and D.W.C. conceived the ideas and designed the methodology; S.B., R.E.C., S.B. and C.L.C. collected the data; R.E.C., S.B., C.L.C and D.W.C. analyzed the data; all authors contributed critically to the drafts and gave final approval for publication.

