## Supporting information for "Plant responses to multiple antagonists are mediated by order of attack and phytohormone crosstalk"

Table S1. MANOVA table for all genes. *Pillai* index shows whether the overall response of all genes was impacted by treatments, while subsequent tables (*F-*statistics) show the response of specific genes.

| MANOVA for all genes |  |  |  |
| --- | --- | --- | --- |
| Fixed Effects | *Pillai* | DF | P-value |
| Aphid Type (A) | 0.942 | 1 | 0.002 |
| Weevil Timing (W) | 1.105 | 2 | 0.348 |
| A ✕ W | 1.509 | 2 | 0.021 |
| Response: PR1 |  |  |  |
| Fixed Effects | *F* | DF | P-value |
| Aphid Type (A) | 25.34 | 1 | < 0.001 |
| Weevil Timing (W) | 4.568 | 2 | 0.033 |
| A ✕ W | 1.617 | 2 | 0.238 |
| Response: GA2ox |  |  |  |
| Fixed Effects | *F* | DF | P-value |
| Aphid Type (A) | 15.09 | 1 | 0.002 |
| Weevil Timing (W) | 1.094 | 2 | 0.365 |
| A ✕ W | 2.571 | 2 | 0.117 |
| Response: AO3 |  |  |  |
| Fixed Effects | *F* | DF | P-value |
| Aphid Type (A) | 5.841 | 1 | 0.032 |
| Weevil Timing (W) | 0.657 | 2 | 0.535 |
| A ✕ W | 1.423 | 2 | 0.278 |
| Response: LOX2 |  |  |  |
| Fixed Effects | *F* | DF | P-value |
| Aphid Type (A) | 7.838 | 1 | 0.016 |
| Weevil Timing (W) | 41.60 | 2 | < 0.001 |
| A ✕ W | 1.505 | 2 | 0.261 |
| Response: ICS1 |  |  |  |
| Fixed Effects | *F* | DF | P-value |
| Aphid Type (A) | 15.54 | 1 | 0.001 |
| Weevil Timing (W) | 0.334 | 2 | 0.722 |
| A ✕ W | 2.473 | 2 | 0.126 |
| Response: DDR230 |  |  |  |
| Fixed Effects | *F* | DF | P-value |
| Aphid Type (A) | 23.32 | 1 | < 0.001 |
| Weevil Timing (W) | 2.903 | 2 | 0.093 |
| A ✕ W | 11.74 | 2 | 0.001 |
| Response: PsLectin |  |  |  |
| Fixed Effects | *F* | DF | P-value |
| Aphid Type (A) | 34.39 | 1 | < 0.001 |
| Weevil Timing (W) | 0.784 | 2 | 0.478 |
| A ✕ W | 2.646 | 2 | 0.111 |

Table S2. MANOVA table for all hormones. *Pillai* index shows whether the overall response of hormones (SA, JA, ABA) was impacted by treatments, while subsequent tables (*F-*statistics) show the response of SA, JA, and ABA.

| MANOVA for all hormones |  |  |  |
| --- | --- | --- | --- |
| Fixed Effects | *Pillai* | DF | P-value |
| Aphid Type (A) | 0.947 | 1 | < 0.001 |
| Weevil Timing (W) | 1.195 | 2 | < 0.001 |
| A ✕ W | 0.877 | 2 | 0.008 |
| Response: Salicylic Acid |  |  |  |
| Fixed Effects | *F* | DF | P-value |
| Aphid Type (A) | 254.2 | 1 | < 0.001 |
| Weevil Timing (W) | 13.25 | 2 | < 0.001 |
| A ✕ W | 7.719 | 2 | 0.006 |
| Response: Jasmonic Acid |  |  |  |
| Fixed Effects | *F* | DF | P-value |
| Aphid Type (A) | 0.966 | 1 | 0.341 |
| Weevil Timing (W) | 5.304 | 2 | 0.018 |
| A ✕ W | 2.653 | 2 | 0.103 |
| Response: Abscisic acid |  |  |  |
| Fixed Effects | *F* | DF | P-value |
| Aphid Type (A) | 9.96 | 1 | 0.006 |
| Weevil Timing (W) | 4.101 | 2 | 0.037 |
| A ✕ W | 4.322 | 2 | 0.032 |

Fig. S1. Non-metric multi-dimensional scaling (NMDS) ordination plot for amino acids analyzed in Fig 5. Amino acids clustered near origin (0,0) had more similar concentrations among the weevil and/or aphid treatments. Amino acids further from the origin (Ala, Lys, Gly, Arg) had less similar concentrations among weevil and/or aphid treatments.
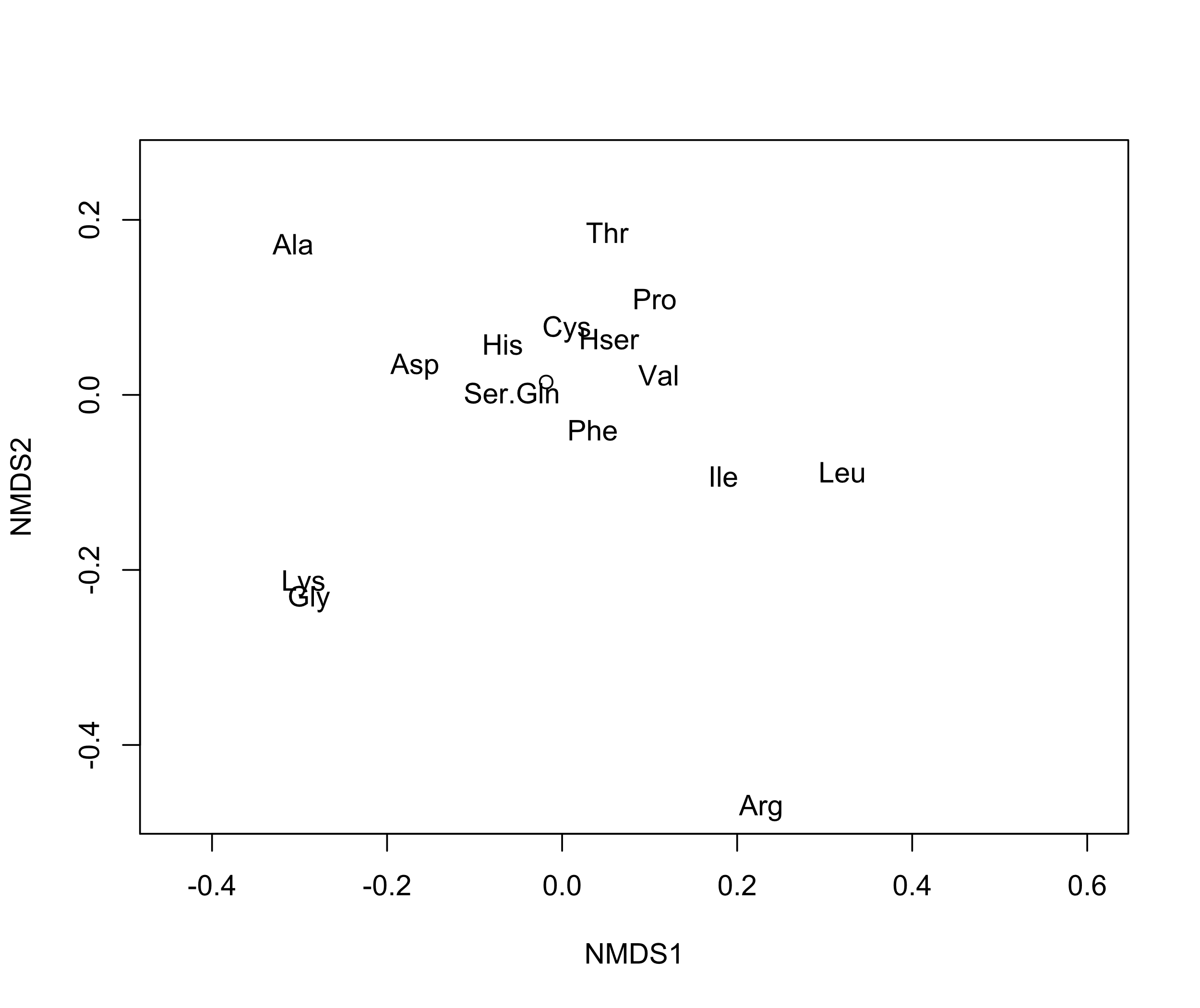

Table S3: List of primers used for this study

| **Gene** | **Primer sequences** | **NCBI Accession No.** | **Amplicon size (bp)** |
| --- | --- | --- | --- |
| *PsPR1 F*  *PsPR1 R* | TGGGGCAGTGGTGACATAAC  TGCGCCAAACAACCTGAGTA | LT635896 | 178 |
| *ICS1 F*  *ICS1 R* | CTGGAACAGGAATAGTGGAAGG  TCAGAGGCAAGTCCAGTTTG | Psat7g237240.1 | 105 |
| *Lox2 F*  *Lox2 R* | GCAACCAAGTGACGAAGTCTA  GGAGACCCGATTGTAAGGTATTT | PsCam 059875 | 95 |
| *PsAO3 F*  *PsAO3 R* | TTATAGGACACAGGCTAGCTCAGCA  TGACACAAGCTTATTCAGCATGACA | EF491600.1 | 127 |
| *PsGA2ox F*  *PsGA2ox R* | GCTGCCACTTAATATTGGAGGATC  GAGTGTTGATGCAAAAGGGGAA | AF101383.1 | 250 |
| *PsDRR230 F*  *PsDRR230 R* | TTGCAGGAACAACGAGCACTT  GCACCAGCAGCGAAAATCAT | AJ308155.1 | 61 |
| *PsLectin F*  *PsLectin* *R* | ATGGGATCCAAGCAACAGAG  CACCATTCTGCAACTTCCAC | EU825771.1 | 92 |
| *PsPOX11* F  *PsPOX11* R | CTTGGAGGACCCACATGGAT  TTTGGCTTGCTGTTCTTGCA | [AB193816.1](https://www.ncbi.nlm.nih.gov/nucleotide/AB193816.1?report=genbank&log$=nuclalign&blast_rank=1&RID=WYJ7ZWJG015) | 61 |
| *β-tubulin* FP  *β-tubulin* RP | GTAACCCAAGCTTTGGTGATC  ACTGAGAGTCCTGTACTGCT | X54844.1 | 203 |
